# The Dietary Fermentable Fiber Inulin Alters the Intestinal Microbiome and Improves Chronic Kidney Disease Mineral-Bone Disorder in a Rat Model of CKD

**DOI:** 10.1101/2023.01.29.526093

**Authors:** Annabel Biruete, Neal X. Chen, Corinne E. Metzger, Shruthi Srinivasan, Kalisha O’Neill, Paul B. Fallen, Austin Fonseca, Hannah E. Wilson, Henriette de Loor, Pieter Evenepoel, Kelly S. Swanson, Matthew R. Allen, Sharon M. Moe

**Author notes:** **Corresponding Author:** Sharon M. Moe, MD, Stuart A. Kleit Distinguished Professor of Medicine, Associate Dean for Clinical and Translational Research, Director, Division of Nephrology and Hypertension, Indiana University School of Medicine, 950 W. Walnut Street, R2-202 Indianapolis, IN 46202.

## Abstract

**Background:** Dietary fiber is important for a healthy diet, but intake is low in CKD patients and the impact this has on the manifestations of CKD-Mineral Bone Disorder (MBD) is unknown.

**Methods:** The Cy/+ rat with progressive CKD was fed a casein-based diet of 0.7% phosphate with 10% inulin (fermentable fiber) or cellulose (non-fermentable fiber) from 22 weeks to either 30 or 32 weeks of age (~30 and ~15 % of normal kidney function). We assessed CKD-MBD, cecal microbiota, and serum gut-derived uremic toxins. Two-way ANOVA was used to evaluate the effect of age and inulin diet, and their interaction.

**Results:** In CKD animals, dietary inulin led to changes in microbiota alpha and beta diversity at 30 and 32 weeks, with higher relative abundance of several taxa, including *Bifidobacterium* and *Bacteroides*, and lower *Lactobacillus*. Inulin reduced serum levels of gut-derived uremic toxins, phosphate, and parathyroid hormone, but not fibroblast growth factor-23. Dietary inulin decreased aorta and cardiac calcification and reduced left ventricular mass index and cardiac fibrosis. Bone turnover and cortical bone parameters were improved with inulin; however, bone mechanical properties were not altered.

**Conclusions:** The addition of the fermentable fiber inulin to the diet of CKD rats led to changes in the gut microbiota composition, lowered gut-derived uremic toxins, and improved most parameters of CKD-MBD. Future studies should assess this fiber as an additive therapy to other pharmacologic and diet interventions in CKD.

**Significance Statement:** Dietary fiber has well established beneficial health effects. However, the impact of fermentable dietary fiber on the intestinal microbiome and CKD-MBD is poorly understood. We used an animal model of progressive CKD and demonstrated that the addition of 10% of the fermentable fiber inulin to the diet altered the intestinal microbiota and lowered circulating gut-derived uremic toxins, phosphorus, and parathyroid hormone. These changes were associated with improved cortical bone parameters, lower vascular calcification, and reduced cardiac hypertrophy, fibrosis and calcification. Taken together, dietary fermentable fiber may be a novel additive intervention to traditional therapies of CKD-MBD.

## Introduction

Chronic kidney disease-mineral bone disorder (CKD-MBD) is a triad of abnormal biochemistries, impaired bone health (renal osteodystrophy), and extra-osseous calcification that results in increased fractures and cardiovascular disease^1, 2^. The incidence of bone fracture is progressively increased at every stage of CKD compared to a similarly aged general population^3^, and the mortality after a hip fracture is doubled in patients with CKD^4^. However, neither bone density nor bone volume fully accounts for the high fracture incidence suggesting CKD specific alterations^5, 6^. Similarly, the risk of cardiovascular events and mortality is increased in CKD and traditional Framingham risk factors and laboratory based diagnostic cardiovascular biomarkers do not fully explain or predict the increased risk^7, 8^. Even aggressive correction of hypertension, anemia and volume overload lead to regression of left ventricular hypertrophy (LVH) in less than 50% of dialysis patients^9^ supporting the importance of non-traditional risk factors specific to CKD.

Uremic toxins are defined as substances elevated in CKD, associated with disease in patients, and toxicity *in vitro*^10, 11^. This includes CKD-MBD toxins such as elevated phosphate, parathyroid hormone (PTH), and fibroblast growth factor-23 (FGF23) that have been linked with arterial calcification, LVH, and myocardial fibrosis in animal models and patients with CKD ^12–17^. In addition, the gut microbiome also plays a critical role in uremic toxin accumulation due to increased production of toxins and/or reduced renal clearance^10, 18, 19^. Microbially-derived uremic toxins can also accumulate from dysbiosis (altered microbiota) and dietary intake. In CKD, dysbiosis is exacerbated by large pill burden including phosphate binders, frequent antibiotic use, and metabolic acidosis^20^. However, little is known about the effects of the gut microbiome on CKD-MBD.

Dietary fiber has a variety of health benefits, including gut health, and is known to affect the gut microbiome in CKD^21, 22^. The effects of dietary fiber may be unique to its physicochemical properties, primarily fermentability and viscosity^21, 23^. Fermentable fiber, including inulin, is metabolized into short-chain fatty acids by commensal members of the gut microbiota and can alter intraluminal pH, improve mineral solubility, and enhance mineral absorption ^24, 25^. In contrast, viscous fiber slows gastric emptying and may encapsulate minerals and limit their absorption^21^. Fermentable dietary fiber has the added benefit of decreasing the serum levels of the gut-derived uremic toxins indoxyl sulfate and p-cresyl sulfate in CKD and hemodialysis patients, but results have been inconsistent^21, 26–28^. The goal of this study was to test the hypothesis that alteration of the gut microbiota by the administration of the fermentable fiber inulin, will improve manifestations of CKD-MBD in our established progressive CKD rat model.

## Material and Methods

### Experimental design

The Cy/+_IU_ colony of rats has been bred at Indiana University for nearly 20 years. Cy/+ rats are characterized by an autosomal dominant progressive cystic kidney disease, but not orthologous to human polycystic kidney disease with gene defects in ciliary proteins. The Cy/+ animals have a mutation in *Anks6*, a gene that codes for the protein SamCystin that binds with *Anks3* and *Bicc1* and thus may alter the nephronophthisis complex^29, 30^. In this rat model, CKD-MBD develops spontaneously, with a much faster progression to end-stage disease in male animals versus females, even with the addition of ovariectomy with a much faster progression to end-stage disease in male animals by 30 to 40 weeks of age versus females, including after ovariectomy ^31, 32^ and therefore only males were used for the current studies. Male Cy/+_IU_ rats (CKD) and normal littermates (NL) were fed the autoclaved grain-based diet (Envigo Teklad 2018SX, Indianapolis IN) until 22 weeks of age and then switched to a casein-based diet (Envigo Teklad TD.04539; 18% casein protein, 0.7% Pi, 0.6% Ca) with cellulose, a minimally fermented fiber, in order to produce a more consistent CKD-MBD phenotype^33, 34^. Treatment began at 22 weeks of age (~ 50% normal GFR). Two concurrent studies (30 week and 32 week duration studies) were conducted and three groups of animals were compared (*n* = 12-14 per group) in each study: 1) Normal littermate animals (NL) with the casein diet with cellulose; 2) CKD animals with the casein diet with cellulose; 3) CKD animals treated with casein diet with inulin (CKD/inulin) instead of cellulose. At 30 weeks of age (~30% normal GFR, study 1) or 32 weeks of age (~15% normal GFR, study 2), animals were anesthetized with isoflurane and underwent cardiac puncture for blood collection followed by exsanguination and bilateral pneumothorax to ensure death. A subset of animals received an injection of calcein (30 mg/kg) fourteen and four days prior to euthanasia. Heart, aorta, kidney, tibia, femora, and intestines were collected, weighed as appropriate, and stored for analyses. Cecal digesta was collected and stored for gut microbiota sequencing. All procedures conducted during this study were approved by the Indianapolis Institutional Animal Care and Use Committee prior to initiating the study.

### Blood Measures

Plasma was analyzed for blood urea nitrogen, creatinine, calcium, and phosphorus using colorimetric assays (Pointe Scientific, Canton, MI, or BioAssay Systems, Hayword, CA). Plasma intact PTH, serum C-terminal and intact FGF23 were determined by ELISA kits according to manufactures instruction (Quidel, San Diego, CA). Serum levels of the oxidative stress marker 8-hydroxy-2’-deoxyguanosine (8-OHdG) were measured using an ELISA kit (Enzo Life Sciences, Farmingdale, NY)^35^. Serum uremic toxins were measured in the lab of Dr. Peter Evenepoel via ultra-performance liquid chromatography-tandem mass spectrometry^36^.

### Gut microbiota of cecal digesta

Rats are cecal fermenters, and therefore total genomic DNA from cecal digesta (proximal large intestine) were extracted (Qiagen Powerlyzer Powersoil kits, Ann Arbor, MI), double-stranded DNA was quantified using the Clariostar spectrometer (BMG Labtech, Cary NC), and quality was assessed by electrophoresis with 2% Agarose EX-gels using the E-Gel iBase (Invitrogen, Grand Island, NY). Fluidigm Access Array was used to generate 16S rRNA gene amplicons, in combination with Roche High Fidelity Fast Start Kit. Primers 515 F (5′- GTGYCAGCMGCCGCGGTAA-3′) and 806R (5′-GGACTACNVGGGTWTCTAAT-3′) targeting a 252 bp-fragment of the V4 region of the bacterial 16S rRNA were amplified ^37^. CS1 forward tag and CS2 reverse tag were added according to the Fluidigm protocol. Sequencing was performed through Illumina Mi-seq using V3 reagents and performed at the Roy J. Carver Biotechnology Center at the University of Illinois. Relative changes in bacterial diversity (α-diversity and β-diversity) and taxonomical changes were analyzed through the open software QIIME2 (version 2019.10^38, 39^). In short, forward reads were imported using the single-end Earth Microbiome Project (EMP) protocol and demultiplexed using the demux plugin. Sequences were then denoised using DADA-2^40^. For diversity analyses, samples were rarified to a sampling depth of 29,050. Amplicon sequence variants (ASVs) were classified using the GreenGenes database version 13_8^41^. A phylogenetic tree was created using the align-to-tree-mafft-fasttree pipeline. The α- and β-diversity metrics were computed using the q2-diversity plugin (core-metrics-phylogenetic) using rarified counts. For α-diversity (number of ASVs, Shannon diversity index, and Faith’s phylogenetic diversity), we tested the difference between CKD with and without inulin at 30 and 32 weeks via 2-way ANOVA, representing the mean of normal rats at 30 and 32 weeks visually in graphs. For β-diversity, we tested the difference between normal, CKD, and CKD with inulin at 30 and 32 weeks using permutational multivariate analysis of variance (PERMANOVA) on unweighted and weighted UniFrac distance matrices and were visualized using EMPEROR.

### Intestinal phosphate transporters, RNA isolation and real-time PCR

Intestinal tissues were flushed with 0.9% saline using a gavage needle to remove luminal contents, cut transversally with scissors, the mucosa scraped with microscope cover slip, and the tissue placed in a microcentrifuge tube and flash-frozen with liquid nitrogen and kept at −80°C until RNA extraction. Duodenum was considered 1 cm proximal to the pylorus until the suspensory muscle of the duodenum; ileum was considered 30 cm proximal to the ileocecal valve, and jejunum the rest of the tissue. Cecum RNA was extracted from whole tissue instead of the mucosa scraping. Total RNA was isolated using the miRNeasy Mini Kit (Qiagen). Target-specific PCR primers were obtained from ThermoFisher Life Technologies: NaPi2b (Rn00584515_m1), Pit1 (Rn00579811_m1), and Pit2 (Rn00568130_m1). The gene expression of NaPi2b, Pit1 and Pit2 were analyzed to assess isoform specificity by real time PCR with Taqman gene expression assays (TaqMan Gene Expression Assays, FAM dye-labeled, Applied Biosystems, Foster City, CA) using ViiA 7 systems as previously described^42, 43^. The ΔΔCT method was used to analyze the relative change in gene expression normalized to *Rplp0*.

### Aortic arch and heart calcification and left ventricular mass index (LVMI)

Segments of the aortic arch and heart were separately washed in 0.9% saline and incubated in 0.6 N HCl for 48 hours. The supernatant was analyzed for calcium using the *o*-cresolphthalein complex 1 method (Calcium kit; Pointe Scientific) and normalized by tissue dry weight as previously described^44, 45^. LVMI was determined by dividing total heart weight by body weight. Heart tissue mRNA expression of transforming growth factor-β (TGF-β) expression was determined by real time qPCR as detailed above with primer Rn00572010_m1.

### Heart Histologic analysis

The hearts were sectioned just below the valves and fixed in 10% (v/v) neutral buffered formalin and paraffin embedded in the Indiana University School of Medicine Histological Core lab. The sections were stained for fibrosis by Masson’s Trichrome stain and calcification by Von Kossa with fast green as previously published ^46^. Bright-field mosaic images of the entire tissue sections were acquired with a Keyence BZ-X810 microscope using a Nikon PlanFluor 10x/0.3 objective lens.

### Bone assessments

Micro-Computed Tomography (microCT) was performed on the proximal tibia using a Skyscan 1172 (Bruker, Billerica, MA) at 12-micron resolution using methods previously published^47–49^. Briefly, a 1 mm region of interest starting roughly 0.5 mm from the distal end of the growth plate was used for analysis of trabecular bone. Cortical bone was assessed as the average of 5 slices 4 mm distal to the trabecular region. All CT analyses were done in accordance with standard guidelines^50^.

For dynamic bone histomorphometry, unmineralized proximal tibia were fixed in neutral buffered formalin then subjected to serial dehydration and embedded in methyl methacrylate (Sigma Aldrich, St. Louis, MO). Serial frontal sections were cut 4 μm-thick and left unstained for analysis of fluorochrome calcein labels. Histomorphometric analyses were performed using BIOQUANT (BIOQUANT Image Analysis, Nashville, TN). A standard region of interest of trabecular bone excluding primary spongiosa and endocortical surfaces was utilized. Total bone surface (BS), single-labeled surface (sLS), double-labeled surface (dLS), and interlabel distances were measured at 20x magnification. Mineralized surface to bone surface (MS/BS; [dLS+sLS/2]/BS*100), mineral apposition rate (MAR; average interlabel distance/10 days) and bone formation rate (BFR/BS; [MS/BS*MAR]*3.65) were calculated. Additional 4-μm sections were stained for tartrate resistant alkaline phosphatase (TRAP) and counter stained with toluidine blue for analysis of osteoclast-covered trabecular surfaces normalized to total trabecular bone surface (Oc.S/BS, %). All nomenclature for histomorphometry follows standard usage^51^.

Prior to mechanical testing, whole femora frozen in phosphate buffered saline were microCT scanned using a Skyscan 1176 (Bruker, Billerica, MA) at 18-micron resolution to assess geometric properties at the mid-diaphysis. Femoral diaphysis mechanical properties were assessed in 3-point bending using standard instrumentation (Test Resources). Bones were thawed to room temperature, kept hydrated in saline, and placed posterior surface down on bottom supports (span = 18 mm). The upper support was brought down in contact with the specimen’s anterior surface and testing was conducted at a displacement rate of 2 mm/min with a 667.2 N load cell. Force versus displacement data were collected at 10 Hz and structural parameters were determined from curves using a customized MATLAB program. Material properties were estimated using standard beam-bending equations^52^.

### Statistics

The question of interest was a comparison between CKD with and without inulin. We have previously shown dramatic differences between CKD and NL littermates, and therefore the NL rat data was used to determine if inulin normalized findings and not included in the statistical analyses (but represented by the black dashed line in all figures). Statistical analyses were conducted by first excluding outliers using ROUT (Q = 1%), followed by a normality test (*p* < 0.05 with Anderson-Darling Test). Data was log-transformed if non-normal before analyses. For comparisons between the two studies (30 and 32 week end points) we used a two-way ANOVA for age (30 vs. 32 weeks) and treatment (inulin vs no inulin) and followed by Tukey’s multiple comparisons test. For end points only assessed at 32 weeks (intestinal phosphate transporter gene expression, bone mechanical test and heart TGF-β expression) we used a one-way ANOVA and, if overall ANOVA showed *p* < 0.05, we conducted within group comparisons by Dunnett’s post hoc analyses, comparing untreated CKD rats versus each of the other groups. The results are expressed as means ± standard deviation (SD) with *p* < 0.05 considered significant (GraphPad Prism Software version 9.4.1, La Jolla, CA).

## Results

### Dietary Inulin lowered the concentration of gut-derived uremic toxins

The CKD serum levels of the uremic toxins indoxyl sulfate, p-cresyl sulfate, p-cresyl glucuronide, phenyl glucuronide, trimethylamine N-oxide (TMAO), and hippuric acid increased from 30 to 32 weeks of age (**Figure 1**, all p< 0.03). Inulin treatment significantly decreased the uremic toxins p-cresyl glucuronide, phenyl glucuronide, and TMAO in CKD rats with an interaction of age and inulin and post-hoc analyses showing inulin lowered levels only at 32 weeks (all p < 0.002). In contrast inulin increased hippuric acid, driven by changes at 30 weeks. Other measured amino acids and their metabolites were not affected by inulin (**Supplemental Figure 1**).

**Figure 1:**
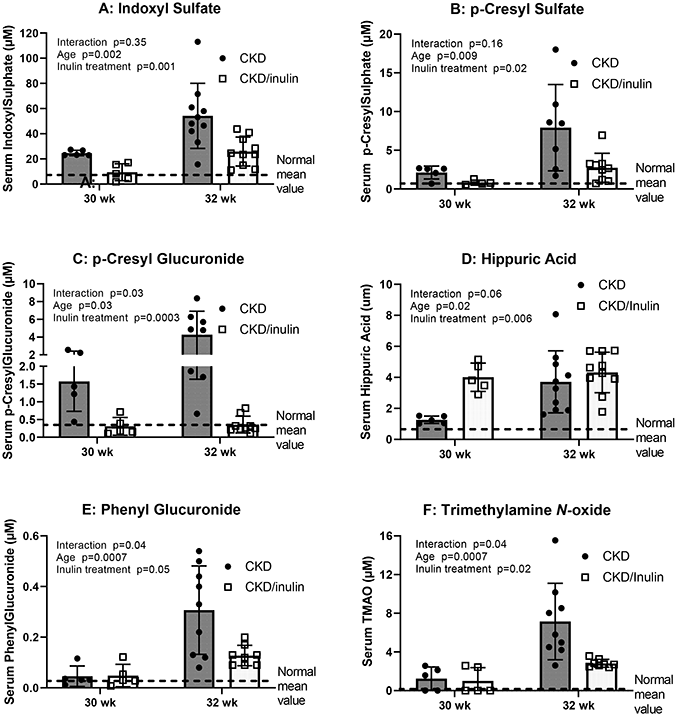
Serum uremic toxin levels are progressively increased with advancing CKD and improved with dietary inulin: Serum was collected at the time of euthanasia and analyzed by mass spectroscopy. Two-way ANOVA compared severity of CKD/age (30 and 32 weeks) and with 10% inulin in the diet (gray bar) compared to cellulose in the diet (black bar). The mean value from the normal animals from both time points are shown as the dashed black line and was not included in the statistical model. The p values for age, inulin, and an interaction of age and inulin are shown in each graph. All CKD measures increased from 30 to 32 weeks. There was an overall effect of inulin for all the toxins, but by post hoc testing the decreases were only at 32 weeks (all p < 0.002). The exception was hippuric acid which showed an increase by inulin at 30 weeks (p = 0.006) and no effect at 32 weeks. N = 5 to 10 for each group shown as individual symbols. (see Supplemental Figure 1 for other uremic toxins for which there was no improvement with inulin).

### Dietary Inulin altered the cecal microbiota

NL and CKD rats had similar α-diversity metrics [Shannon index (**Figure 2A**), ASVs, and Faith’s phylogenetic diversity (**Supplemental Figure 2**)].CKD rats fed inulin had lower α-diversity at 30 weeks (p<0.0008), but similar to NL and CKD at 32 weeks (p>0.72 **Figure 2A**). β-diversity, or diversity between samples, using unweighted (presence vs. absence) and weighted (taking abundance into consideration) UniFrac distances also showed that the overall microbial composition was similar between NL and CKD at 30 and 32 weeks (PERMANOVA q>0.05), but CKD rats fed inulin was different from both at both time points (PERMANOVA q<0.01, **Supplemental Figure 2**).

**Figure 2:**
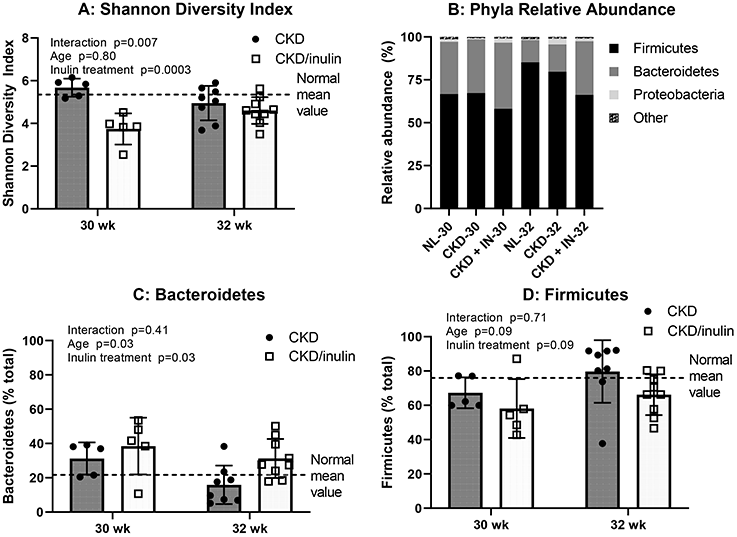
Inulin altered the cecal microbiota: Cecal genomic DNA was extracted and the V4 region of the 16S rRNA gene was sequenced and analyzed at 30 and 32 weeks. Two-way ANOVA compared severity of CKD/age (30 and 32 weeks) and with 10% inulin in the diet (gray bar) compared to cellulose in the diet (black bar). The mean value from the normal animals from both time points are shown as the dashed black line and was not included in the statistical model. The p values for age, inulin, and an interaction of age and inulin are shown in each graph. Panel A shows that the Shannon diversity index, a metric of α-diversity, was lower with inulin at 30 weeks (*post hoc* p=0.0008), but there was no difference at 32 weeks. Panel B shows the average relative abundance at the phyla-level in each group, where the major phyla were Firmicutes and Bacteroidetes accounting for over 95% of the relative abundance in all groups. Panel C and D shows a main effect of inulin increasing the relative abundance of Bacteroidetes and a trend toward decreasing Firmicutes.

Taxonomical analyses at the phyla-level showed that Firmicutes and Bacteroidetes were the most abundant phyla across all rat groups accounting for over 95% of the relative abundance in all groups at both time points. Phyla relative abundance was similar between NL and CKD, but inulin-treated rats had higher Bacteroidetes (p=0.03 **Figure 2B and 2C**) and tended to have lower the relative abundance of Firmicutes (p=0.09, **Figure 2D)**. At the genus-level, composition was similar between NL and CKD. When comparing CKD with and without inulin, the inulin treatment led to a higher relative abundance of *Bifidobacterium, Bacteroides, Sutterella, Bernesiellaceae, Allobaculum*, S24-7, and unclassified Lachnospiraceae (all p<0.001), and a lower relative abundance of *Lactobacillus, Oscillospira, Adlercreutzia, Dorea*, and unclassified Clostridiaceae, Ruminococcaceae, Rikenellaceae, and Peptostreptococcaceae (all p<0.03, **Figure 3, Supplemental Figure 3**).

**Figure 3:**
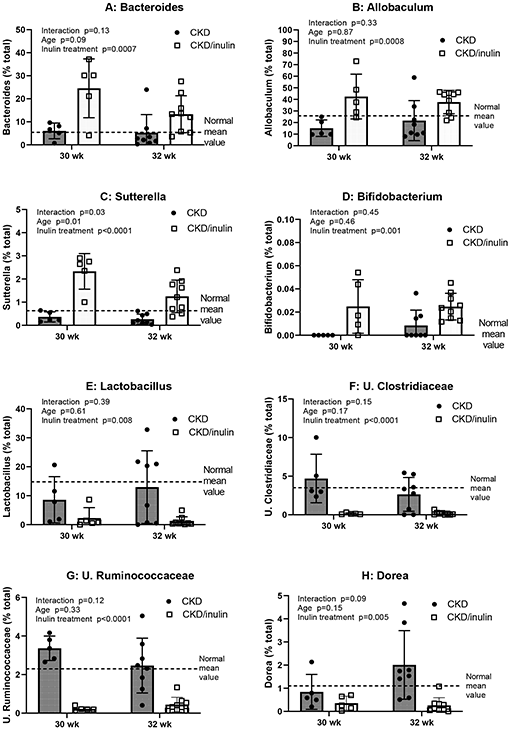
Inulin treatment led to taxonomical changes in the cecal microbiota. Cecal genomic DNA was extracted and the V4 region of the 16S rRNA gene was sequenced and analyzed at 30 and 32 weeks. Two-way ANOVA compared severity of CKD/age (30 and 32 weeks) and with 10% inulin in the diet (gray bar) compared to cellulose in the diet (black bar). The mean value from the normal animals from both time points are shown as the dashed black line and was not included in the statistical model. The p values for age, inulin, and an interaction of age and inulin are shown in each graph. Panels A-D shows the genera that increased after inulin treatment (*Bacteroides, Allobaculum, Sutterella*, and *Bifidobacterium)* all of them showing a main effect of inulin and only *Sutterella* with an inulin-by-age interaction, where inulin-treatment had a higher relative abundance compared to untreated CKD. Panels E-H shows the genera in which inulin treatment lowered relative abundance (*Lactobacillus*, unclassified Clostridiaceae and Ruminococcaceae, and *Dorea)*, all of them showing a main effect of inulin. Data from normal littermates is shown as the dashed black line and was not included in the statistical model. The p values for age, inulin, and an interaction of age and inulin are shown in each graph. There was no change from 30 to 32 weeks in any of the genera.

### Dietary Inulin did not alter CKD progression, but improved some CKD-MBD biochemistries

CKD rats had a progressive decline in kidney function (elevated BUN, creatinine) and increase in kidney weight due to cyst growth from 30 to 32 weeks, both different than NL animals. Inulin had no effect on kidney function (**Supplemental Table 1**). The plasma phosphorus (**Figure 4A**, p = 0.01) and PTH (**Figure 4B**, p = 0.0004) levels increased in CKD rats from 30 to 32 weeks. Inulin treatment significantly decreased both plasma phosphorus and PTH levels driven by changes at 32 weeks **(Figure 4A and 4B; post hoc PTH at 32 weeks p = 0.0004)**. There was a decrease in calcium levels from 30 to 32 weeks, but no effect of inulin **(Figure 4C)**. Both c-terminal FGF23 and intact FGF23 increased from 30 to 32 weeks, with an effect of inulin to increase intact FGF23, driven by 30 week data **(Figure 4D** and **Figure 4E)**. Oxidative stress as measured by 8-OHdG, a marker of DNA oxidation, did not increase over time, with a modest reduction by inulin (p = 0.04; F**igure 4F**), driven by 32 week results. Overall, these results demonstrated that inulin did not alter CKD progression but reduced plasma phosphate and PTH. We therefore examined if there was a direct effect of inulin on intestinal phosphate transporters in each intestinal segment at 32 weeks demonstrating no effect on the expression of NaPi2b, Pit1, or Pit2 **(Supplemental Figure 4)**.

**Figure 4:**
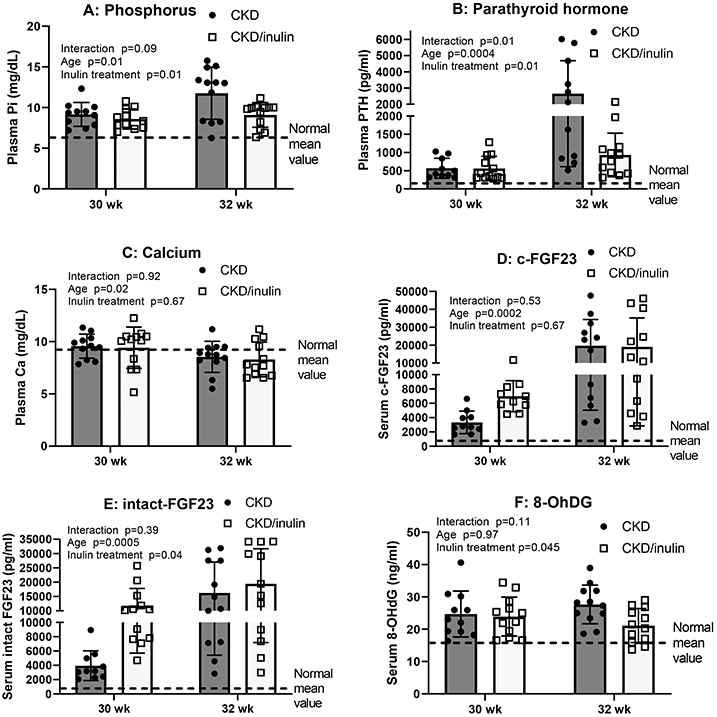
Biochemistries of CKD-MBD progressively increased with advancing CKD with variable effect of dietary inulin: Serum/plasma was collected at the time of euthanasia and analyzed by either colorimetric assays or ELISA. Two-way ANOVA compared severity of CKD/age (30 and 32 weeks) and with 10% inulin in the diet (gray bar) compared to cellulose in the diet (black bar). The mean value from the normal animals from both time points are shown as the dashed black line and was not included in the statistical model. The p values for age, inulin, and an interaction of age and inulin are shown in each graph. All measures increased from 30 to 32 weeks except calcium and 8-OHdG, a measure of oxidative stress. At 32 weeks, post hoc testing showed that inulin significantly lowered phosphorus, PTH and 8-OHdG (all p < 0.01). At 30 weeks, inulin increased iFGF23 (p = 0.048), but not at 32 weeks. N = 10 to 12 for each group shown as individual symbols.

### Dietary Inulin treatment improved cardiovascular parameters in CKD rats

We have previously demonstrated spontaneous cardiac and vascular calcification in our CKD model at older ages^34, 53^, both of which contribute to LVH and arrythmias^54^. We replicated these findings in the current study, demonstrating increased aorta calcification with progression from 30 to 32 weeks (p = 0.001; **Figure 5A**) and a reduction by inulin overall (p = 0.003). Cardiac calcification did not significantly increase from 30 to 32 weeks (p = 0.11), with a trend towards an effect of inulin overall (p = 0.07; **Figure 5B**); post hoc testing showing significance at 32 weeks (p = 0.003). Left ventricular mass index (LVMI) increased from 30 to 32 weeks (p = 0.005) and decreased with inulin (p = 0.049; **Figure 5C**). We therefore assessed the heart mRNA expression of TGF-β at 32 weeks of age and found increased expression in the CKD animals with a reduction by inulin (**Figure 5D**). Qualitative histological evaluation showed the calcification was primarily in the arterioles with surrounding fibrosis **(Supplemental Figure 5**).

**Figure 5:**
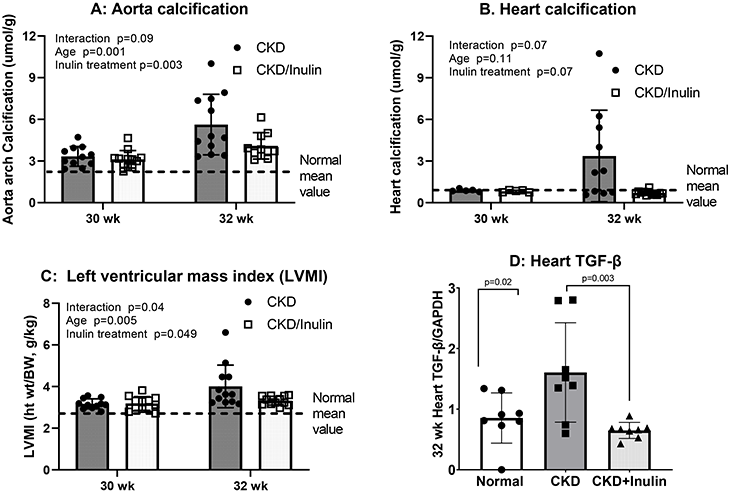
Inulin reduced aorta and cardiac calcification and decreased transforming growth factor-β expression: The heart was collected after euthanasia, weighed to calculate left ventricular mass index (LVMI = heart weight/body weight; C) and then the ascending aorta and part of the ventricle collected for the quantification of calcification (A and B). Two-way ANOVA compared severity of CKD/age (30 and 32 weeks) and with 10% inulin in the diet (gray bar) compared to cellulose in the diet (black bar). The mean value from the normal animals from both time points are shown as the dashed black line and was not included in the statistical model. The p values for age, inulin, and an interaction of age and inulin are shown in each graph. Aorta calcification increased from 30 to 32 weeks, and was decreased by inulin at 32 weeks by post hoc testing (p = 0.009; Panel A). There was an overall trend towards an effect of inulin (p = 0.07, Panel B) with heart calcification, but only at 32 weeks. Left ventricular mass index (Panel C) increased from 30 to 32 weeks, with a trend towards a reduction by inulin (overall p = 0.049) but only at 32 weeks (p = 0.01 by post hoc testing). With these latter changes, we investigated the mRNA expression of TGF-β in the left ventricle at 32 weeks, and observed an increase with CKD compared to NL and a reduction with inulin (Panel D). N = 7 to 10 for each group shown as individual symbols.

### The effect of inulin treatment on bone in CKD rats

MicroCT analysis of cortical bone at the proximal tibia showed cortical porosity (**Figure 6A**) dramatically increased from 30 to 32 weeks in CKD rats (p = 0.01) and was reduced by inulin (p = 0.035); post hoc testing showed inulin’s effect only at 32 weeks (p = 0.001). Cortical area and thickness (**Figures 6B and 6C**) decreased from 30 to 32 weeks (p = 0.0003, p = 0.0005, respectively) and both improved with inulin treatment (p =0.023, p = 0.002, respectively). These changes were both driven by changes at 32 weeks (both p < 0.007 by post hoc testing). In contrast, there was no age effect or inulin effect on trabecular bone volume (**Figure 6D**) or trabecular thickness (not shown). There was no age effect or inulin effect of the bone volume/total volume (**Figure 6D**) or trabecular thickness (not shown). There was no age effect on trabecular separation and number (**Figure 6 E and 6F**), but inulin decreased trabecular separation and increased trabecular number (p = 0.009 and 0.04, respectively).

**Figure 6:**
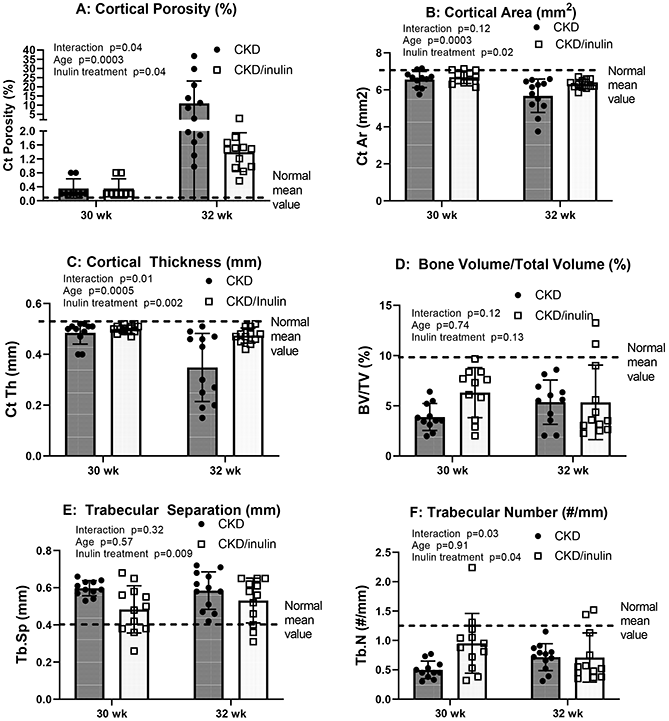
Inulin treatment improved cortical bone changes assessed by microCT, with some effects on trabecular bone: Bones were collected at the time of euthanasia and analyzed by microCT. Two-way ANOVA compared severity of CKD/age (30 and 32 weeks) and with 10% inulin in the diet (gray bar) compared to cellulose in the diet (black bar). The mean value from the normal animals from both time points are shown as the dashed black line and was not included in the statistical model. The p values for age, inulin, and an interaction of age and inulin are shown in each graph. Cortical porosity (A) increased with age, and was improved with inulin in the diet and interaction between these. Cortical area (B) and thickness (C) decreased from 30 to 32 weeks and was improved with inulin treatment, but only at 32 weeks by post hoc testing (p < 0.007 for both measures). In contrast, there was no age effect or inulin effect on trabecular volume/total volume (**D**). There was no age effect on trabecular separation and number (**E and F**), but inulin decreased trabecular separation and increased trabecular number (p = 0.009 and 0.04, respectively). N = 10-12 for each group shown as individual symbols.

Dynamic histomorphometry demonstrated higher trabecular bone formation rate (BFR/BS) and mineral apposition rate (MAR) in CKD rats at both 30 and 32 weeks compared to normal (**Figure 7A-B**), but no increase with age. Inulin treatment decreased bone formation rate (p = 0.028) with a similar non-significant trend for MAR (p = 0.06). In contrast, there was decreased mineralizing surface (MS/BS) with age (p = 0.015) but no effect of inulin (**Figure 7C**). Trabecular bone osteoclast surface (OcS/BS; **Figure 7D**) progressively increased from 30 to 32 weeks (p = 0.0002). Inulin decreased osteoclast surface (p = 0.0002), with post hoc comparison only different at 32 weeks (p < 0.0001)

**Figure 7:**
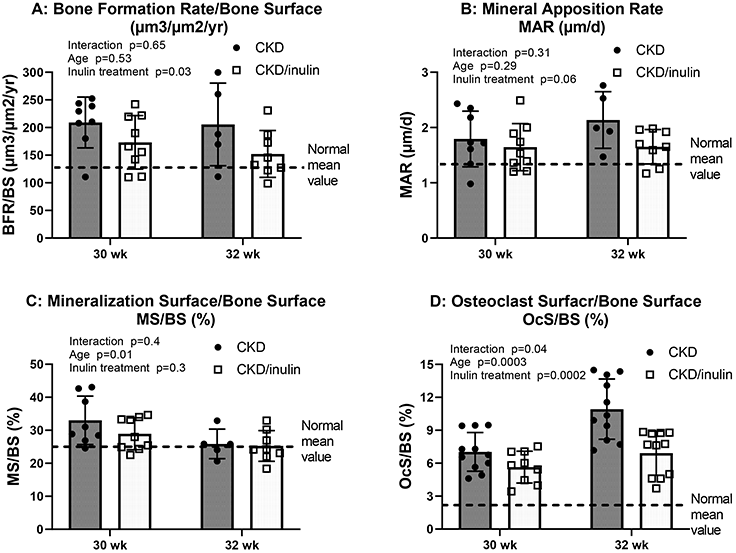
Inulin decreased CKD induced high bone turnover: Bones were collected at the time of euthanasia and embedded for histomorphometry. Two-way ANOVA compared severity of CKD/age (30 and 32 weeks) and with 10% inulin in the diet (gray bar) compared to cellulose in the diet (black bar). The mean value from the normal animals from both time points are shown as the dashed black line and was not included in the statistical model. The p values for age, inulin, and an interaction of age and inulin are shown in each graph. Bone formation rate (BFR/BS; A) and mineral apposition rate (MAR; B) in CKD rats did not increase with advancing age/CKD; inulin treatment decreased bone formation rate (p = 0.028) with a similar trend for MAR (p = 0.06). In contrast, mineralizing surface (MS/BS; C) decreased with age (p = 0.015) but was unaffected by inulin. Bone tartrate resistant alkaline phosphatase (TRAP) staining to quantify osteoclast surface (OcS/BS; **D**) progressively increased from 30 to 32 weeks (p = 0.0002). Inulin decreased osteoclast surface (p = 0.0002), with post hoc comparison only different at 32 weeks (p < 0.0001) N = 5-12 for each group shown as individual symbols.

Mechanical properties of the femur analyzed only at 32 weeks, demonstrated clear differences between CKD and NL animals for ultimate load, post-yield displacement, stiffness, total work, ultimate stress, and total toughness (all p < 0.01), but no effect of inulin treatment on any parameter **(Supplemental Figure 6)**.

## Discussion

In this study we examined the impact on the administration of the fermentable fiber inulin on CKD-MBD. We examined two time points, 30 and 32 weeks in the Cy/+ rat, where there are dramatic and parallel changes observed in patients with CKD early stage 4 versus 5 in CKD-MBD. It is this transition where the increased PTH and FGF23 moves from appropriate compensation for decreased renal excretion of phosphate to a state where the persistent increase of PTH and FGF23 can no longer compensate, and end-organ effects of CKD-MBD such as bone loss due to cortical porosity, vascular calcification, and LVH ensue^55, 56^. Our results showed marked increases in PTH, FGF23, aorta calcification, cortical porosity, and uremic toxins between 30 and 32 weeks. The administration of inulin altered the gut microbiota and dramatically decreased gut-derived uremic toxins including indoxyl sulfate and p-cresyl sulfate at both ages, and reduced the rise in phosphorus and PTH, and extra-skeletal calcification between 30 and 32 weeks. In contrast, inulin had little effect on FGF23 levels but there was still a reduction in LVH and aorta/cardiac calcification. The increased bone remodeling (bone formation rate, osteoclast surface) and cortical changes (porosity, thickness) were also reduced by inulin, effectively preventing the dramatic changes between 30 and 32 weeks. However, we did not see inulin-induced alterations in trabecular bone volume or bone mechanics indicating inulin impact some, but not all measures of bone. Thus, inulin had positive effects, but did not completely reverse the sequelae of CKD-MBD suggesting additive therapies such as calcimimetics or vitamin D may be needed.

Our study demonstrated a reduction in serum phosphate and PTH, although levels remained above the normal range. The mechanism for these effects are unknown. We did not see an effect of inulin on intestinal phosphate transporters, albeit we only measured mRNA expression and not protein expression or flux and did not assess colonic transporter expression. Alternatively, inulin may have affected bone remodeling through reductions in indoxyl sulfate. Inulin increased FGF23 at 30 weeks but had no effect at 32 weeks for unclear reasons. We did find increased FGF23 mRNA expression in total bone at 30 weeks confirming increased production (unpublished observation). It is conceivable that the inulin induced increase in FGF23 at 30 weeks led to a reduction in PTH at 32 weeks with enhanced urinary phosphate excretion. However, a previous study in unilateral nephrectomy rats given partially hydrolyzed guar gum (minimally viscous, fermentable fiber) and found a decrease in urinary phosphate excretion^57^, going against this potential explanation. We did not see a change in serum calcium to account for the changes in PTH, although we measured total calcium. Additional studies that assess ionized calcium in the blood and urinary excretion of both ions are needed to further understand this complex physiology. Another hypothesis is that changes were due to reduced food intake by our animals, but there was no change in weight. Unraveling this will require balance studies with controlled feeding in our rat model.

The addition of inulin to the diet altered the cecal microbiota alpha and beta diversity, and important taxonomical changes, including a higher relative abundance of *Allobaculum, Bifidobacterium, Bacteroides*, and unclassified Lachnospiraceae. This translated to a reduction in most commonly studied gut-derived uremic toxins, and we postulate that effect was the main driver of the positive effects on bone and vascular calcification. Two meta analyses of prebiotic, probiotic, and synbiotic therapies identified conflicting results on whether these interventions lowered indoxyl sulfate, but both studies (similar to our study) reported decreased p-cresol sulfate levels with interventions^58, 59^. Indoxyl sulfate is a ligand for the aryl hydrocarbon receptor (AhR) that is critical in the removal of both environmental and endogenous toxins ^60^. AhR null mice have impaired bone formation and decreased osteoclast differentiation as assessed by reduced receptor activator of factor kappa-B ligand (RANKL)^61^. At levels comparable to patients with CKD, *in vitro* studies of indoxyl sulfate suppress mineralization in rodent osteoblasts^62^, downregulate the PTH receptor expression in primary mouse osteoblasts^63^ and inhibit RANKL-dependent differentiation of osteoclasts^64, 65^. In young mice with CKD from 5/6th nephrectomy, indoxyl sulfate worsened bone quantity by microCT^66^. Indole fed to rats after a parathyroidectomy was additive in decreasing bone formation by histomorphometry and alkaline phosphatase activity to the low PTH^67^. These results are consistent with our findings of reduced bone formation rate, decreased osteoclast surfaces, and decreased cortical porosity, although it is not possible to separate out primary effects of inulin from secondary effects from the decrease in PTH. Unfortunately, as inulin did not positively impact bone mechanical properties, it likely that additional interventions will be necessary. Indeed, despite the “reduced” PTH with inulin at 32 weeks, levels were still over 1000 pg/ml, so it is likely concomitant therapy with PTH lowering agents will be needed to normalize bone.

In the current study, inulin reduced arterial calcification and trended to reduce cardiac calcification and left ventricular mass index. We have previously demonstrated that this rat model develops concentric LVH with fibrosis (with increased expression of transforming growth factor beta; TGF-β) and associated arrhythmias^54^. It is interesting that we also observed that inulin reduced TGF-β indicating that inulin may improve cardiac outcomes in CKD, the leading cause of death. Elevated phosphorus levels are traditionally thought of the key mediator of arterial calcification^68^. However, in rodents, the oral administration of indoxyl sulfate led to arterial calcification in the adenine rat, supporting *in vitro* studies showing a direct effect this uremic toxin to induce calcification in cultured vascular smooth muscle cells^69^. Thus, it is plausible that a reduction in phosphorus and a reduction in indoxyl sulfate may be additive to reduce arterial calcification.

Despite these exciting findings in our animal model, the importance of gut-derived uremic toxins in humans is not clear, with inconsistent results. A study of 1,170 Japanese dialysis patients reported an association of indoxyl sulfate levels with all-cause mortality but no association with cardiovascular events^70^. In China, a study of 258 hemodialysis patients reported levels of indoxyl sulfate above the median was associated with a hazard ratio of 5.31 (95% CI 2.43-11.58) for heart failure events, but as a continuous variable levels were not associated with outcomes. A large US hemodialysis study of 1,273 participants found no relationship with indoxyl sulfate levels and adjudicated cardiovascular outcomes^71^. However, in non-dialysis advanced CKD a study of 139 patients reported that indoxyl sulfate was associated with all-cause mortality, cardiovascular mortality, and increased coronary artery calcification^72^. An intestinal spherical carbon toxin adsorbent, AST-120, that lowers many uremic toxins including indoxyl sulfate, did not lead to a reduction in the progression of CKD in patients in randomized trials^73^, and therefore was not FDA approved. However, a recent study in mice with ischemia reperfusion-induced acute kidney injury treated with AST-120 reported improvements in cardiac dysfunction by echocardiography^74^. A second study in dogs with induced heart failure showed AST-120 treatment was shown to reduce both myocardial apoptosis and fibrosis along with decreases in TGF-β1 expression^75^. These studies suggest that intestinal toxin reduction with fiber and/or absorbents may prove beneficial to cardiac dysfunction in CKD, but much work remains to confirm this exciting possibility.

In conclusion, we observed that the administration of inulin to the diet improved parameters of CKD-MBD in our preclinical model. While the presumed beneficial effect was a reduction in gut-derived uremic toxins, additional mechanistic studies are warranted.

## Author Contributions

SMM, AB, NXC, and MA conceived and designed the study. AB, NXC, CEM, SS, KO, PF, AF, HEW, HD, PE, KSS performed the experiments; AB, NCX, SMM, MA, analyzed and interpreted the results. SMM, AB, NCX drafted the manuscript and all authors approved the final version of the manuscript.

## Acknowledgements

None

## Disclosures

There is no related conflicts of interest. AB has received honoraria from Amgen. SMM is a scientific consultant for Amgen, Sanifit, and Ardelyx who have other products in the field of CKD-MBD.

## Funding

AB was supported by T32 DK120524-02 and Indiana-CTSI KL2 (with support from Grant Numbers, KL2TR002530 (Sheri Robb, PI), and UL1TR002529 (Sarah Wiehe and Sharon Moe, co-PIs) from the National Institutes of Health, National Center for Advancing Translational Sciences, Clinical and Translational Sciences Award.) AF was funded, in part, with support from the Short-Term Training Program in Biomedical Sciences Grant funded, in part by T35HL110854 from the National Institutes of Health. SMM was supported by NIH UL1TR002529 and P30 AR072581-06.

## Data sharing sources

N/A

## Notes

### Competing Interest Statement

The authors have declared no competing interest.

