## Supplemental Figures for "The Dietary Fermentable Fiber Inulin Alters the Intestinal Microbiome and Improves Chronic Kidney Disease Mineral-Bone Disorder in a Rat Model of CKD"

**SUPPLEMENT:**

**Supplemental Table 1**: Change in kidney function and body weight (additive to Figure 4)

**Supplemental Figure 1**: Other measured metabolites that were not reduced with inulin (additive to Figure 1)

**Supplemental Figure 2:** Inulin treatment leads to changes in α-diversity and β-diversity (additive to Figure 2)

**Supplemental Figure 3:** Additional taxa that were differentially affected by inulin (additive to Figure 3)

**Supplemental Figure 4:** Intestinal phosphate transporter expression at 32 weeks

**Supplemental Figure 5:** Representative Histology of Hearts from Normal, CKD, and CKD animals treated with inulin

**Supplemental Figure 6:** Bone mechanical properties were reduced in CKD and not improved with inulin treatment

### Supplemental Table 1: Change in kidney function and body weight

| Treatment groups | Body weight (g) | | Kidney weight (g) | | BUN (mg/dL) | | Creatinine (mg/dL) | |
|---|---|---|---|---|---|---|---|---|
| | 30 weeks | 32 weeks | 30 weeks | 32 weeks | 30 weeks | 32 weeks | 30 weeks | 32 weeks |
| Normal | 525 ±36 | 524±19 | 3.44±0.38 | 3.45±0.16 | 18.21±2.41 | 17.35±3.34 | 0.59±0.13 | 0.63±0.08 |
| CKD | 503±35 | 453±77* | 5.72±0.71* | 7.44±1.46* | 43.60±5.86* | 46.15±10.12* | 1.27±0.29* | 1.96±0.81* |
| CKD/ 10% inulin | 508±33 | 488±42* | 6.24±1.41* | 6.35±0.88*# | 45.13±7.59* | 42.18±5.33* | 1.22±0.25* | 1.73±0.41* |

Data are shown as mean ± SD (n =10-12 rats each group). *p < 0.05, NL vs. CKD; #p<0.05, CKD vs. CKD/10% inulin by ANOVA at each time point (30 and 32 weeks).

### Supplemental Figure 1: Other measured metabolites that were not reduced by inulin

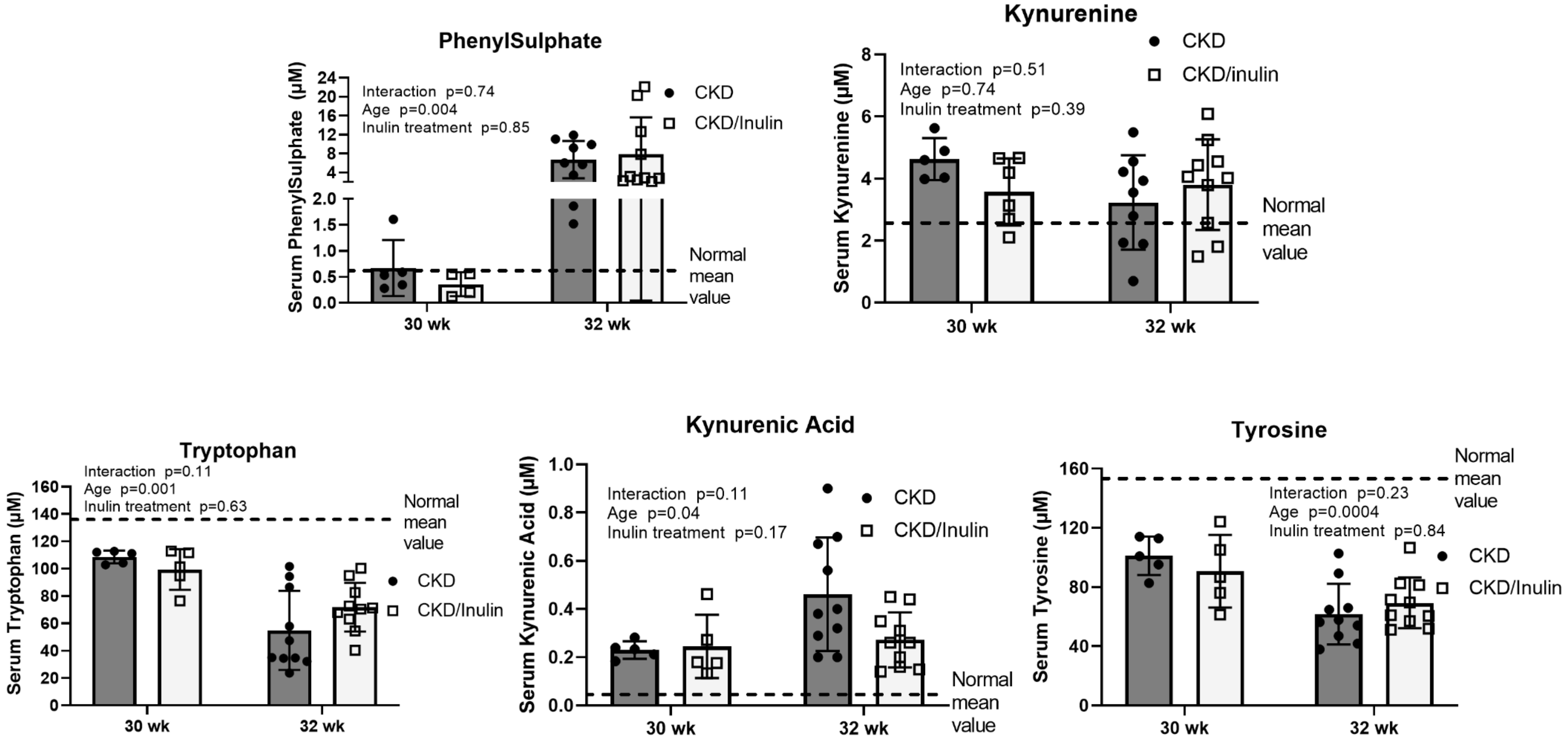


**Supplemental Figure 1: Uremic toxins that were not reduced by inulin:** Serum was collected at the time of euthanasia and analyzed by mass spectroscopy. Two-way ANOVA compared severity of CKD/age (30 and 32 weeks) and with 10% inulin in the diet (white bar) compared to cellulose in the diet. The mean value from the normal animals from both time points are shown as the dashed black line and was not included in the statistical model. The p values for age, inulin, and an interaction of age and inulin are shown in each graph. Except for kynurenine, levels changed with increased age, but inulin had no effect. The p values for age, inulin, and an interaction of age and inulin are shown in each graph. N = 5 -10 for each group.

**Supplemental Figure 2: Inulin treatment leads to changes in α-diversity and β-diversity.**

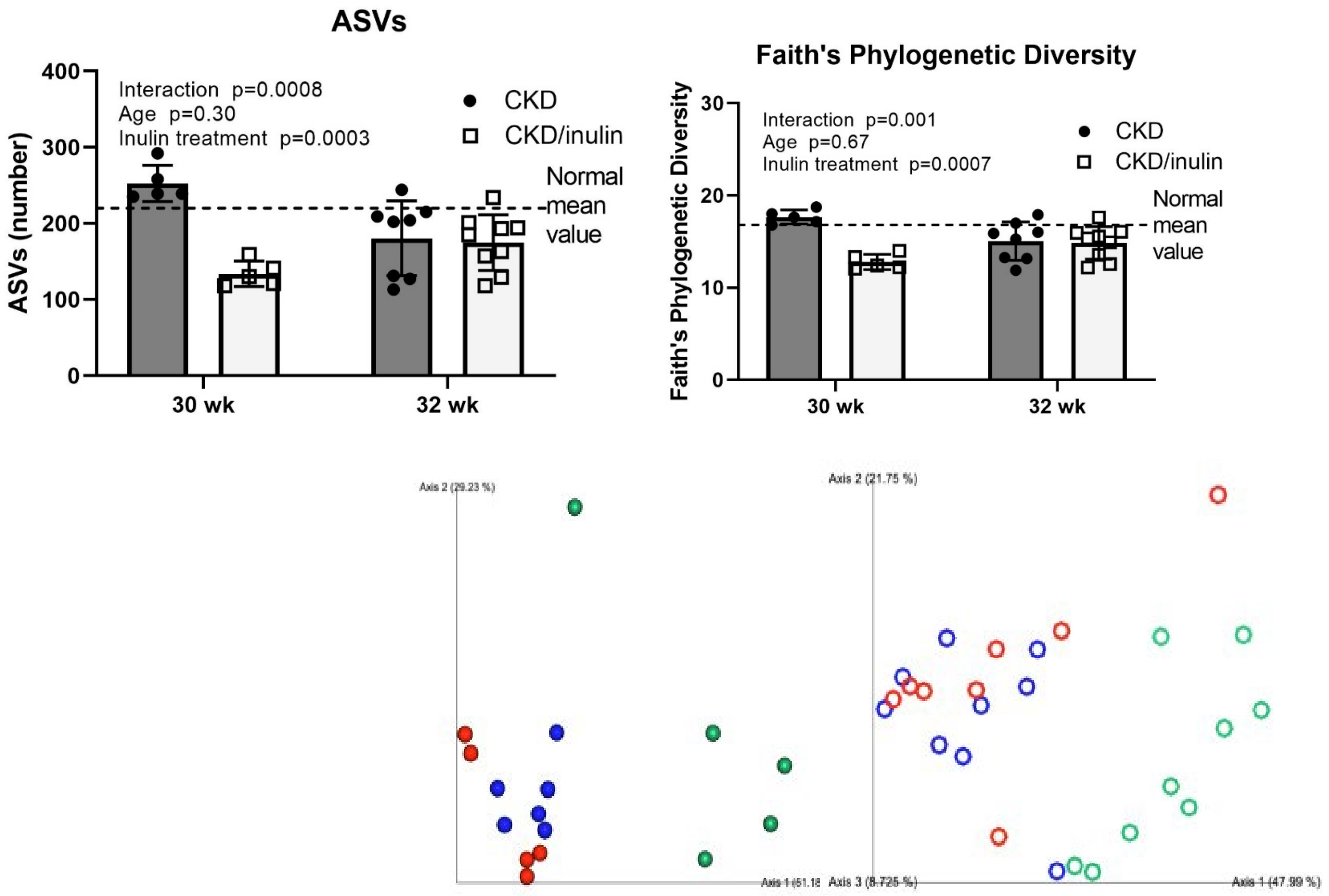


**Supplemental Figure 2:** Inulin treatment leads to changes in α-diversity and β-diversity. Two-way ANOVA compared severity of CKD/age (30 and 32 weeks) and with 10% inulin in the diet (white bar) compared to cellulose in the diet. The mean value from the normal animals from both time points are shown as the dashed black line and was not included in the statistical model. Top left: Amplicon sequence variants (ASVs) were classified using the GreenGenes database version 13_8. Top right: Faith's phylogenetic diversity. The plot below shows the weighted UniFrac distances at 30 (closed circles, NL blue, CKD red, CKD/inulin green) and at 32 weeks (open circles NL blue, CKD red, CKD/inulin green) where the overall microbial composition was similar between NL and CKD rats at 30 and 32 weeks (PERMANOVA q>0.05), but inulin-treated rats were different (PERMANOVA q<0.05). Supplemental Figure 3: Additional taxa that were differentially affected by inulin.

**Supplemental Figure 3: Additional taxa differentially affected by inulin.**

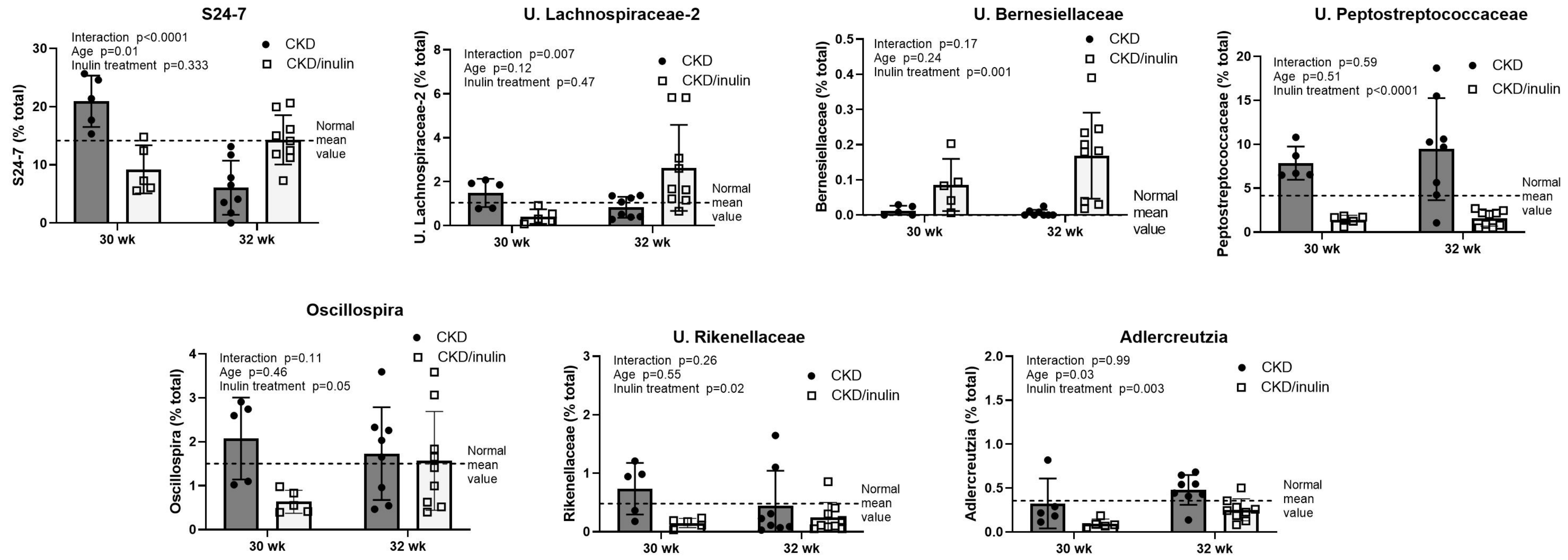


**Supplemental Figure 3:** Two-way ANOVA compared severity of CKD/age (30 and 32 weeks) and with 10% inulin in the diet (white bar) compared to cellulose in the diet. The mean value from the normal animals from both time points are shown as the dashed black line and was not included in the statistical model. Inulin led to initial lower relative abundance of S24-7 and unclassified Lachnospiraceae at 30 weeks, but the abundance increased at 32 weeks. Unclassified Bernesiellaceae had a higher relative abundance at 30 and 32 weeks with inulin. Unclassified Peptostretococcaceae had a lower relative abundance with inulin at 30 and 32 weeks. Oscillospira had an initial lower relative abundance with inulin at 30 weeks, but similar to untreated CKD rats at 32 weeks. Unclassified Rikenellaceae and Adlercreutzia had a lower relative abundance with inulin treatment at both time points.

**Supplement Figure 4: Intestinal expression of phosphate transporters at 32 weeks**

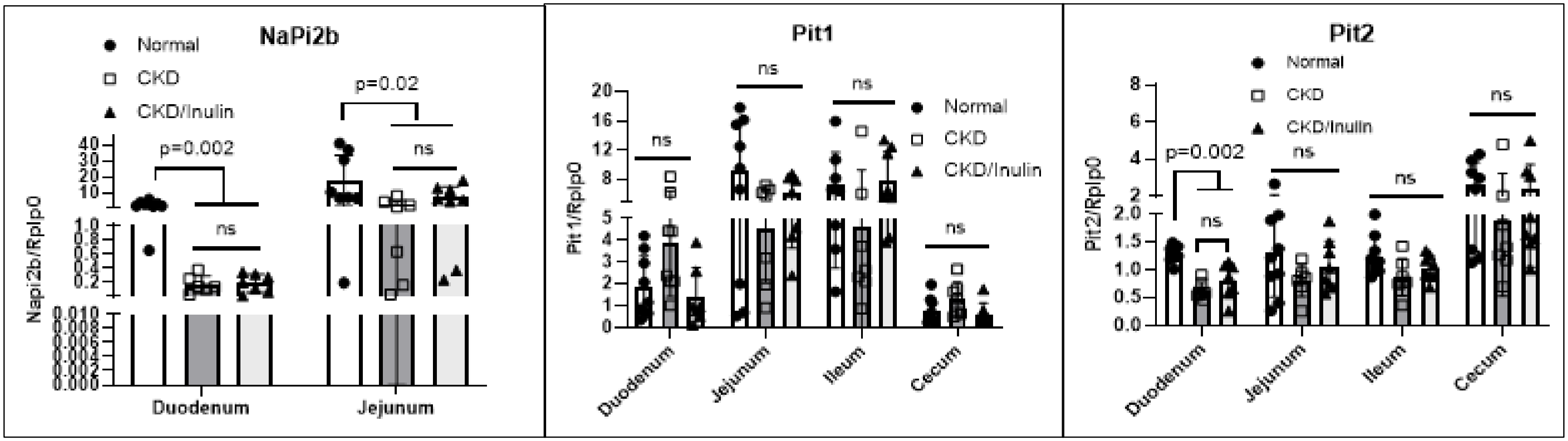


**Supplemental Figure 4: Intestinal expression of phosphate transporters at 32 weeks:** Each intestinal segment was examined by real time PCR for the expression of phosphate transporters and the groups compared by ANOVA. Each graph shows individual transporter expression for different segments for normal animals (black dot symbols/white bars), CKD animals (square symbols/dark gray bars), and CKD + inulin (triangle symbols/light gray bars)transporters. NaPi2b expression was reduced in CKD animals compared to NL in the duodenum and jejunum, but not affected by inulin and not expressed in the remaining segments. Pit1 was not different in any of the segments. Pit2 was reduced in CKD animals compared to NL in the duodenum and not affected by inulin. N = 8 to 10 per group.

**Supplemental Figure 5: Representative Histology of Hearts from Normal, CKD, and CKD animals treated with inulin.**

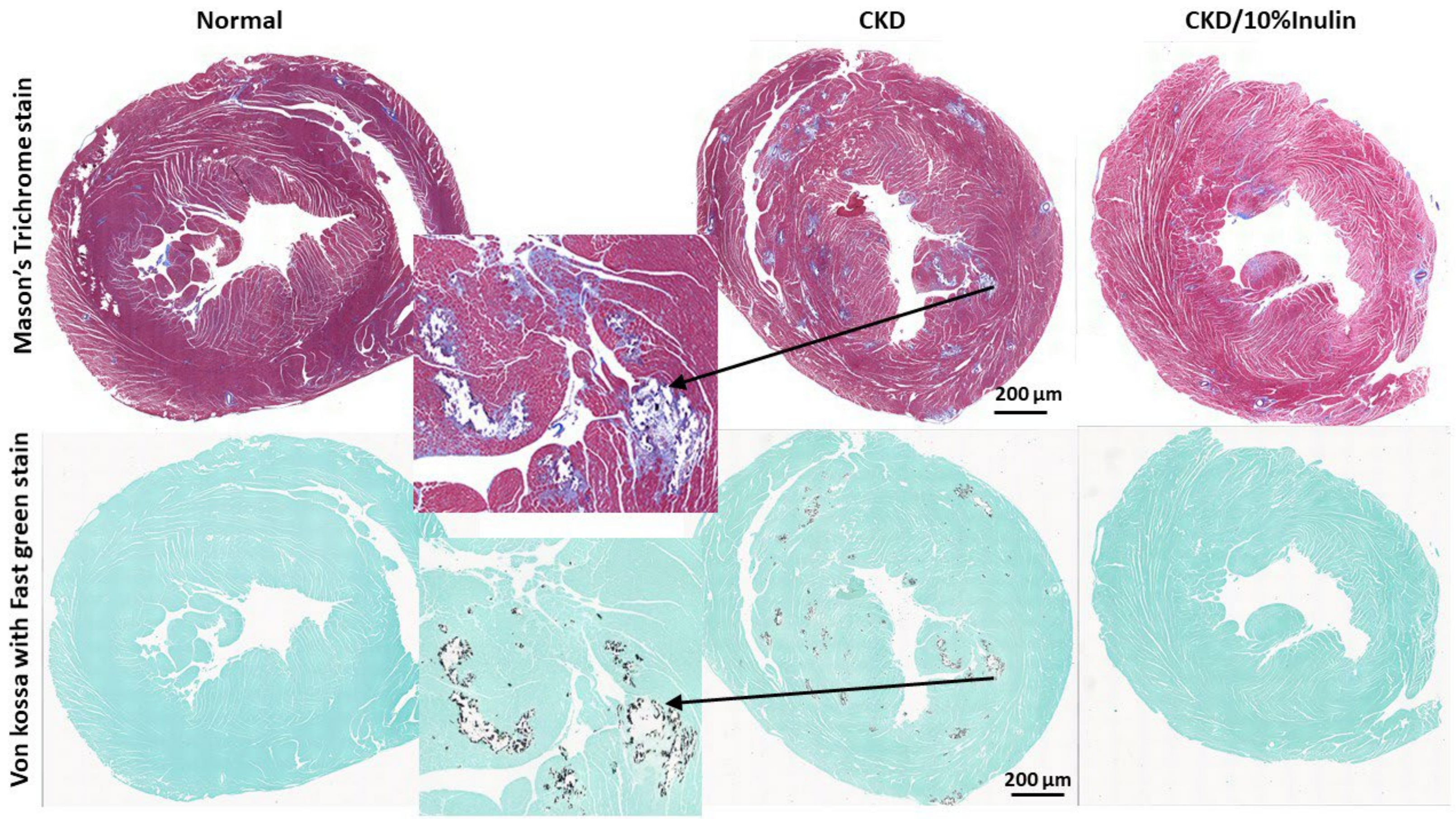


**Supplemental Figure 5:** Representative cross sections taken just below the valves stained for fibrosis with Mason's Trichrome stain (top panel) and calcification by Von Kossa with fast green background staining (bottom panel). The CKD animals (middle panels) had increased fibrosis demonstrated by Mason's Trichrome stain (top insert blue staining) located primarily around the arterioles that were calcified (bottom insert black staining). In contrast, the fibrosis and calcification were reduced in animals given dietary inulin (far right set of panels). These data corroborate the quantification of calcium and TGFβ expression in Figure 5 in the manuscript. The line represents 200 µm.

### Supplemental Figure 6: Bone mechanical properties were reduced in CKD and not improved with inulin treatment

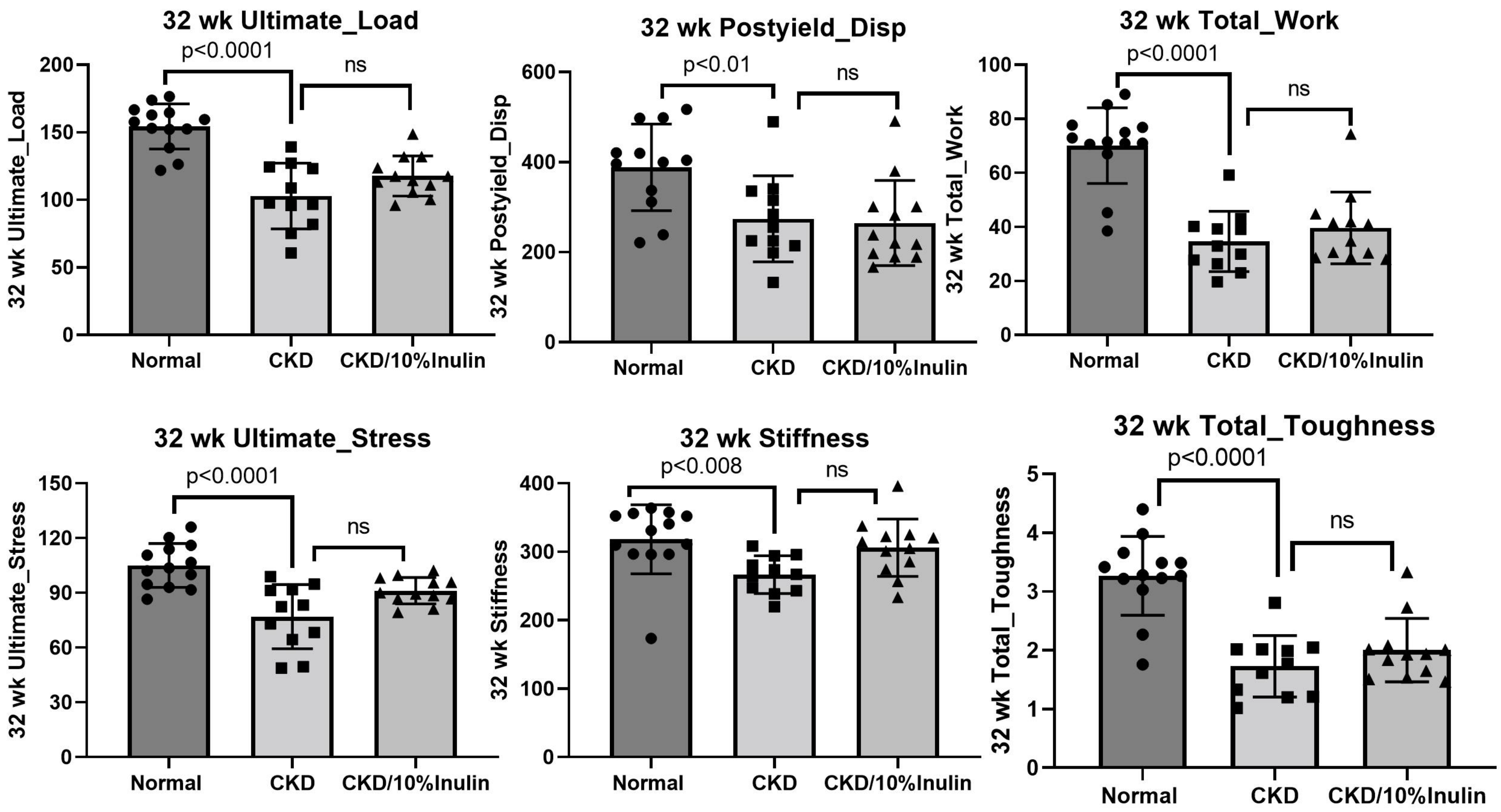


**Supplemental Figure 6**: Bone was collected at the time of euthanasia at 32 weeks and analyzed by 4-point bending. Three groups were compared by ANOVA with post hoc testing:  Normal animals (black circle symbols, dark gray), CKD fed cellulose diet (black square symbols, light gray bar) and CKD animals fed inulin (black triangle symbols, medium gray bar).  As shown in all of the panels, CKD bone quality was significantly worse to NL animals by all measures, but there was no improvement seen when the animals were fed inulin. N = 10-12 for each group shown as individual symbols.
